# Meta-Analysis of ERP Investigations of Pain Empathy underlines methodological issues in ERP research

**DOI:** 10.1101/225474

**Authors:** Michel-Pierre Coll

**Affiliations:** Department of Experimental Psyhology, University of Oxford, Anna Watts Building, Radcliffe Observatory Quarter, Woodstock Road, Oxford OX2

**Keywords:** Empathy, Pain, ERP, Methods, Meta-analysis

## Abstract

Empathy has received considerable attention from the field of cognitive and social neuroscience. A significant portion of these studies used the event-related potential (ERP) technique to study the mechanisms of empathy for pain in others in different conditions and clinical populations. These show that specific ERP components measured during the observation of pain in others are modulated by several factors and altered in clinical populations. However, issues present in this literature such as analytical flexibility and lack of type 1 error control raise doubts regarding the validity and reliability of these conclusions. The current study compiled the results and methodological characteristics of 40 studies using ERP to study empathy of pain in others. The results of the meta-analysis suggest that the centro-parietal P3 and late positive potential component are sensitive to the observation of pain in others, while the early N1 and N2 components are not reliably associated with vicarious pain observation. The review of the methodological characteristics shows that the presence of selective reporting, analytical flexibility and lack of type 1 error control compromise the interpretation of these results. The implication of these results for the study of empathy and potential solutions to improve future investigations are discussed.

## Introduction

Empathy is a complex psychological construct that refers to the ability of individuals to share the experience of others (Batson, 2009; Coll et al., 2017; Cuff et al., 2016). It is of high importance for healthy interactions with others and has been suggested to be altered in several psychiatric conditions (Bird & Viding, 2014; Decety & Moriguchi, 2007). With the hope that understanding the neuronal mechanisms of empathy will bring new insights on this concept, it has been one of the main endeavours of social neuroscience to describe the cerebral processes and computations underlying empathy (de Vignemont & Singer, 2006; Decety & Jackson, 2004; Klimecki & Singer, 2013).

While it is challenging to elicit empathy in a controlled neuroimaging experiment, studies often use cues of nociceptive stimulation in others (i.e. a needle piercing a hand) to study empathy since they are relatively unambiguous, highly salient and easily understood (Vachon-Presseau et al., 2012). In the electroencephalography (EEG) literature, the event-related potential (ERP) technique has mostly been used to study this phenomenon by measuring electrical brain responses to nociceptive cues depicting various levels of pain in others. In a seminal study, Fan and Han, 2008 showed participants real pictures or cartoon depictions of hands in painful or neutral situations and asked participants to either judge the intensity of the pain experienced or to count the number of hands present in the stimuli. The results showed an early effect of pain in the N1 and N2 component that was not influenced by task demands and a later effect of pain in the P3 component that was modulated by task requirements. The authors interpreted these results as the presence of an early automatic response indexing emotional sharing and a late response indexing the cognitive evaluation of others’ pain (Fan and Han, 2008). Similar paradigms are now regularly used to study pain empathy in healthy and clinical samples (see Results section). Although innovative, the study by Fan and Han, 2008 has several limitations. By analysing multiple time-windows at several scalp locations, the authors to perform over 100 statistical tests on ERP data without adjusting the significance threshold for those multiple comparisons. This suggests that some results have a high probability of being false positives (Kilner, 2013) and that the effect of vicarious pain observation on ERP therefore deserves further scrutiny.

The issue of multiple comparisons is a common problem in neuroimaging studies due to the large amount of data collected (Luck & Gaspelin, 2017; Poldrack et al., 2008). In the ERP literature, this is often made worse by the traditional use of factorial analyses performed in several time windows and at several scalp locations without clear hypotheses on the main effects and interactions (Luck & Gaspelin, 2017). Furthermore, this large amount of data also allows for considerable analytical flexibility; that is the idea that the same dataset can be analysed in different ways with significant changes in the results and interpretations depending on the analytical pipeline chosen (Carp, 2012). The presence of flexibility in design and analysis choices and ambiguity regarding how to best to make these choices can lead researchers to compare the results of different analytical pipelines and choose the one which gives the most favourable pattern of result (Carp, 2012; Simmons et al., 2011). When considerable analytical variability is present in a particular field without justification, it can raise doubt regarding the validity of the results and their interpretations.

If the study of pain empathy using ERP is to provide results that are appropriate to further our understanding of empathy in different contexts and populations, it seems imperative to assess 1) the reliability of the effect of the observation of pain in others on the ERP response and 2) the amount of variability and flexibility in the designs employed to investigate this phenomenon. To reach these aims, a review of the methodological practices used in 40 ERP studies investigating pain empathy and a meta-analytical compilation of their results was performed. The results provide meta-analytical evidence for the association between late ERP components and the observation of pain in others. However, there was considerable variation in the design and analyses and incomplete reporting of results, raising doubts on the validity of some of these results.

## Methods

### Study selection

A systematic review of the literature was performed following the Preferred Reporting Items for Systematic Reviews and Meta-Analyses (PRISMA) guidelines (Moher et al., 2009). The articles included in this review were selected by searching *PubMed* for studies that were available online before May 1, 2018 using different combinations of keywords (e.g. “EEG”, “ERP”, “Pain”, “Empathy”, “Vicarious”, see Table S1). The reference lists and citations reports of eligible studies were also consulted.

To be included in this report, studies had to report scalp ERP data in response to pictures depicting nociceptive stimulations (e.g. Jackson et al. (2005)). Studies using facial expressions stimuli were included only when nociceptive stimulations were visible in the stimuli (e.g. needle piercing the skin of the face). This procedure led to the selection of 40 studies published between 2008 and 2018 in 20 different journals (see Table S1 and asterisks in the references list).

From the 40 studies reviewed, 4 were excluded from the quantitative meta-analysis. One was excluded because it used the same dataset and analyses as another study (Han et al., 2008), one because it did not report sufficient information (Ikezawa et al., 2012), one because it used non-parametric statistics (Fitzgibbon et al., 2012) and one because it reported incorrect degrees of freedom and *F* statistics (Sun et al., 2017). The quantitative meta-analysis was therefore performed on 36 studies (marked with double asterisks in the References). The PRISMA flowchart for study selection and rejection is shown in Figure 1. The data reported in this review were manually extracted from the text of the published articles or accompanying supplementary materials and available in Table S1.

**Fig. 1.**
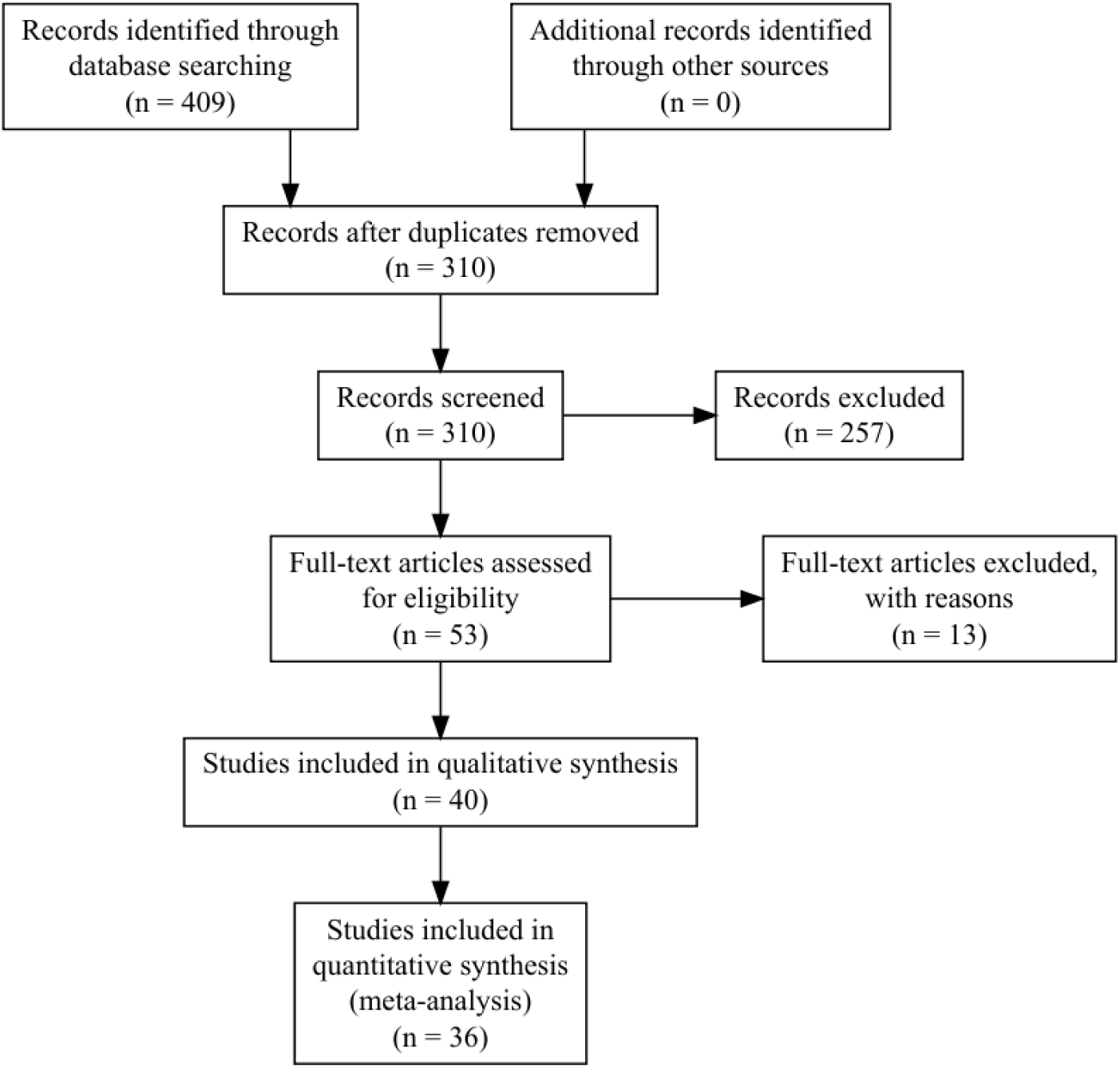
PRISMA flowchart. Reasons for exclusions for studies not included are shown in Table S1.

### Methodological review

Several variables concerning the Participants, Materials and procedures, Data collection and preprocessing, ERP measurements, Statistical analyses and the Reporting of results were collected and summarized below. When information was not clearly reported, the value was estimated based on the available information or assumed to have a particular value (e.g. when no post-hoc correction was reported, it was assumed that none were used). When insufficient information was available for a particular variable, it was marked as not reported and treated as a missing value.

In order to assess the exhaustiveness of the hypotheses formulated regarding ERPs in each study, hypotheses were classified in one of four categories: *Complete, Partial, Alternative* and *None*. Hypotheses were rated as *Complete* if they clearly predicted the specific components that were expected to be influenced by all manipulations as well the direction and the location of this effect. If some predictions were present but were incomplete or unclear, the hypothesis was rated as *Partial*. Hypotheses that were formulated as two alternative outcomes without a clear prediction were labeled *Alternative* and the absence of prediction regarding ERP effects was labeled *None*. This procedure was applied separately for the factorial analysis of variance performed on ERP components and for the correlational analysis of ERP components with other variables.

### Meta-analysis of ERP components

A quantitative meta-analysis was carried out to assess the evidence for a modulation of different ERP components by the observation of pain. However, this was complicated by the fact that most studies reviewed reported only significant results (see Section 3.5 Results reporting) and the general lack of clarity and precision of the results section of many studies. Nevertheless, when possible, *F*-values for the omnibus repeated measures test comparing the ERP response to pain and neutral stimuli were collected for each study.

When several between and within-subject factors were manipulated, the *F*-value of the baseline condition was selected when available. (e.g. in healthy controls or following neutral priming). Similarly, when available, the *F* values for individual electrodes were collected. However, in most cases, the omnibus *F* value from the main effect of pain in a multi-factorial analysis was collected and attributed to all electrodes included in the analysis. When mean amplitudes and standard deviations or standard errors were reported, the paired sample *t*-value for the pain effect was calculated assuming a correlation of 0.70 between measurements. When only the exact p-value was reported, the corresponding *t*-value was found using the t distribution. Following available guidelines (Cooper & Hedges, 1994; Moran et al., 2017), when the effect was reported as non-significant without the information necessary to compute an effect size, the effect size was calculated assuming *p* = 0.5.

This was done for each of the most frequently analysed components (N1, N2, P3, LPP). All *F* values were subsequently converted to *t*-values by taking their square root (Brozek & Alexander, 1950). In order to compare the effect sizes across studies, the *t*-values collected for each electrode and component were converted to Hedges’s *g*, a standardized measure of difference that is less biased than Cohen’s *d*, especially for small samples (Hedges, 1981). Effects were scored as positive when the observation of pain led to increased ERP amplitude (i.e. more positive) than the observation of neutral stimuli, and as negative when the ERP amplitude was more positive in response to neutral stimuli compared to pain stimuli.

Effect sizes were summarised in different ways. First, the spatial distribution of the effects was assessed by plotting scalp maps of the weighted absolute effect size for each component of interest. The absolute effect was taken to show where the effects were stronger on the scalp independently of their direction. The average effect at each electrode was weighted by the number of studies including this electrode in their analysis in order to decrease the weight of the effects at electrodes that were only analysed in a small number of studies. Second, the proportion of significant effects and significant interactions with other factors was compiled for each component. Third, a random effect meta-analysis was performed for each component at fronto-central (Fz, FCz, F1, F2, F3, F4, FC1, FC2, FC3, FC4), centro-parietal (Cz, CPz, C1, C2, C3, C4, CP1, CP2, CP3, CP4) and parieto-occipital (Pz, POz, P1, P2, P3, P4, PO1, PO2, PO3, PO4) electrode clusters to estimate summary effect size and the heterogeneity across studies. Finally, potential publication bias was assessed using funnel plots and regression tests (Egger et al., 1997). To assess potential excess significance, the number of studies finding a significant effect for each component was compared to the expected number of significant studies given the power of each study to detect the summary effect size using exact one-tailed binomial tests and a significance threshold of 0.10 (Ioannidis & Trikalinos, 2007).

### Data Availability

All data and scripts used to produce this manuscript and accompanying figures, the PRISMA guidelines checklist and supplementary information and figures are available online. All data processing and analyses were performed using *Rstudio* (RStudio Team, 2015; R Core Team, 2018) and the Fieldtrip toolbox (Oostenveld et al., 2011) within Matlab R2017a (The MathWorks, Inc., Natick, Massachusetts, United States).

## Systematic review of methodological practices

### Goals and hypotheses

All studies reviewed aimed at comparing the effect of an experimental manipulation and/or participant characteristics on the ERP to pain stimuli with the goal of furthering the understanding of the mechanisms underlying empathy in the general population and/or in various clinical groups. As shown in Table 1, the majority of studies presented incomplete hypotheses regarding the analysis of ERP components. While some studies provided complete hypothesis for the factorial analysis of ERPs, this was rarely the case for correlational analyses of ERP and other behavioural or physiological variables, suggesting that most of these analyses were exploratory in nature.

**Table 1.** Ratings of the the exhaustiveness of hypotheses for the factorial and correlational analyses of ERP data.

| Judgment of hypothesis | Factorial analyses on<br>ERP data (% of 40 studies) | Correlations with<br>ERP data (% of 26 studies) |
| --- | --- | --- |
| None | 10 | 69.23 |
| Partial | 50 | 26.92 |
| Complete | 22.5 | 0 |
| Alternative | 17.5 | 3.85 |

### Participants

In the 40 studies reviewed, 42.5% used a between-subject design and 57.5% used a within-subject design. Among studies employing a between-subjects design, 22.5% compared participants from the general population to participants from a clinical group. These clinical conditions included autism, amputation, bipolar disorder, fibromyalgia, juvenile delinquents and schizophrenia. Only 2 studies provided a justification for their sample size based on *a priori* power analyses. The average sample size, sample size per group and participants excluded are shown in Figure 2. In order to asses the power of each study to detect a small (*d* = 0.2), medium (*d* = 0.5) or large (*d* = 0.8) effect size (Cohen, 1992), a power analysis was performed for each study and each of these effect size using the sample size per group, a two-sided paired t-test and a significance threshold of 0.05. As shown in Figure 2, most studies were only adequately powered to detect a large effect size equal to or higher than d = 0.8. No studies had 80% power to detect a small or medium effect size.

**Fig. 2.**
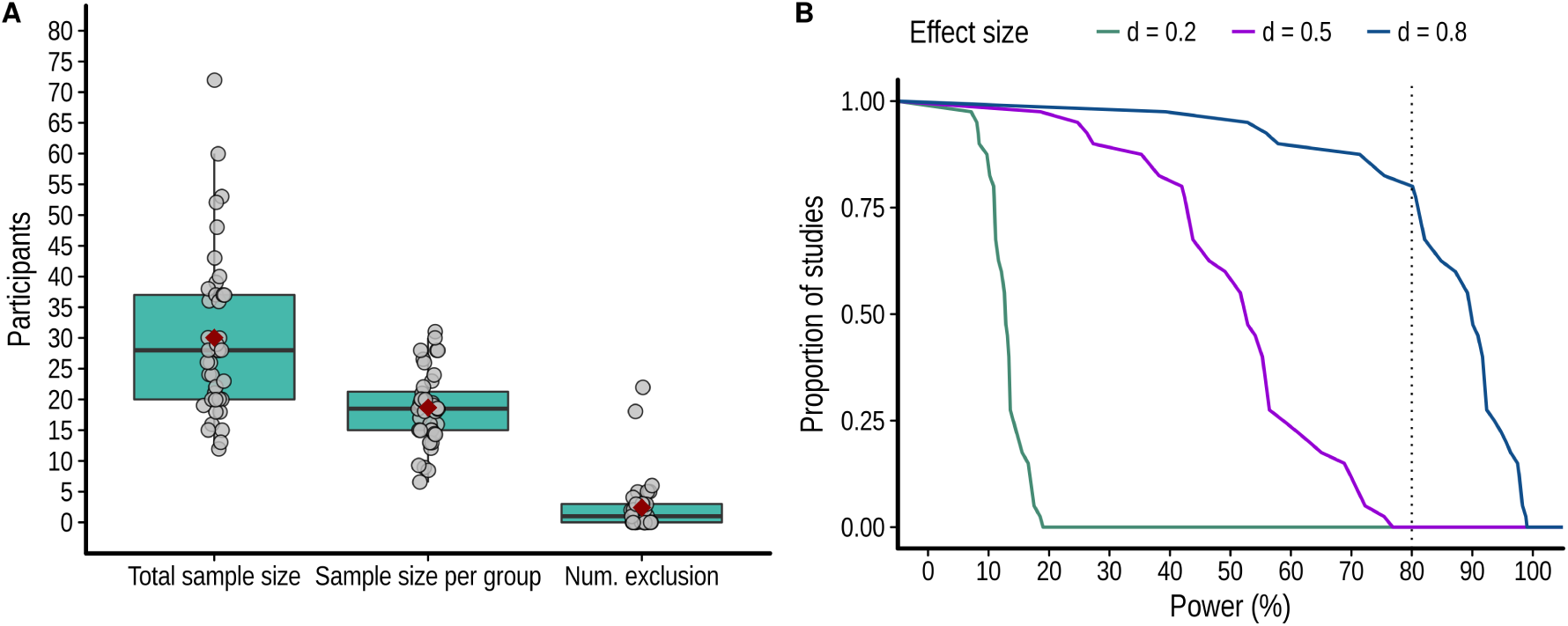
**A** Total sample size, sample size per group and number of exclusion. For all box plots in this paper, the boxes show the 25th and 75th percentile, the error bars show 1.5 interquartile interval, the horizontal line shows the median, the red square shows the mean and the gray points show the individual studies. **B** Proportion of studies as a function of the level of power to detect a small (d = 0.2), medium (d = 0.5) and large (d = 0.8) effect size.

### Materials and procedure

***Visual stimuli***. The majority of studies reviewed (82.5%) used static pictures depicting limbs (hands or feet) in painful and non-painful situations similar to those used initially by Jackson et al. (2005). The other types of stimuli included short 3-frames clips of limbs in painful situations (7.5%), static pictures of faces pricked by a needle or touched with a cotton bud (5%), both faces and limbs (2.5%) and anthropomorphised objects pricked by a needle or touched with a cotton bud (2.5%). The average stimulus duration is shown in Figure 3.

**Fig. 3.**
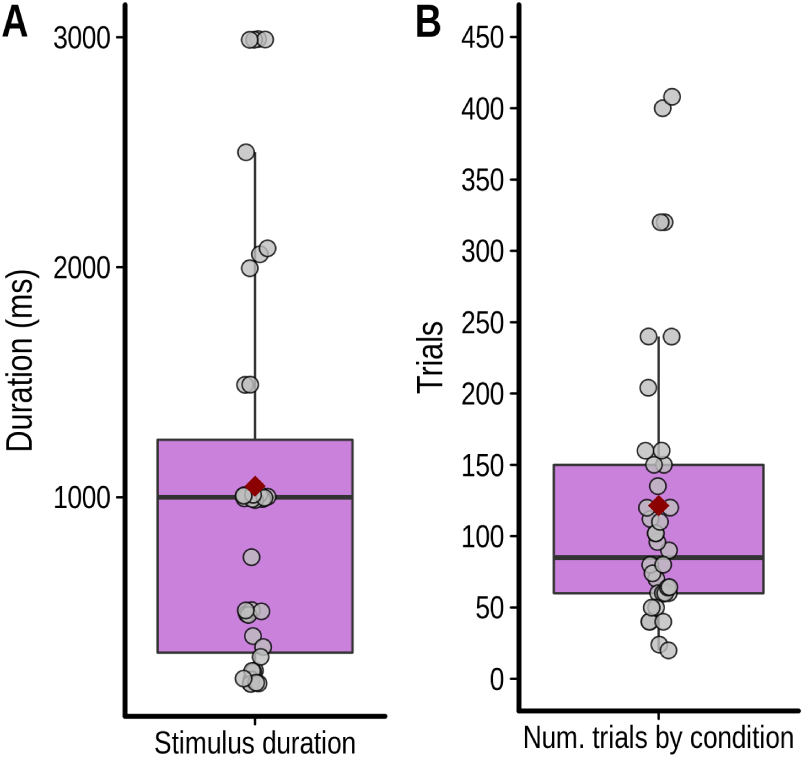
**(A)** Stimulus duration and **(B)** number of trials per condition.

***Experimental task***. All studies compared ERPs to painful and non-painful stimuli. Including this Pain factor, studies had on average 2.08 within-subjects factors (SD = 0.45, range: 1–3) and an average of 4.55 within-subjects conditions (SD = 1.67, range: 2–8). During the experimental tasks, the participants were either asked to detect the presence of pain in a forced choice format (60% of studies), to assess the intensity of the pain observed using a rating scale (17.5%), to passively observe the pictures (15%) or to perform another behavioural task (7.5%). The average number of trials per condition is shown in Figure 3.

***Recordings***. EEG was collected from 60–64 scalp electrodes in the majority of cases (70 %) while the remaining studies used 32 (27.58%), 72 (5%) or 128 (5%) scalp electrodes. 30 % of studies did not report the manufacturer of the EEG system used. The majority of studies (27.5 %) used an EEG system manufactured by *Brain Products*. Other manufacturers included *Biosemi* (27.5 %), *EGI Geodesic* (5%) and *NuAmps* (10%).

***Preprocessing***. EEG data were high-pass filtered in most cases with a cutoff value of 0.1 Hz (52.5%) and the rest of the studies used a high-pass cutoff between 0.01 and 1 Hz. For low-pass filters, the most used cutoffs were 100 Hz and 30 Hz used in respectively 30% and 40% of cases. Other low pass filters cutoffs included values between 40 and 80 Hz. 7.5% and 5% of studies did not report using a high-pass filter or a low-pass filter respectively. 5% of studies reported using a notch filter to filter out electrical noise.

The average of the mastoid processes and the average of all scalp electrodes were the most popular reference schemes for EEG analyses (37.5% and 40%respectively). Other studies use the average of the earlobes (7.5%), a single mastoid (12.5%) or did not report the reference used for analysis (2.5%). EEG data were epoched for analyses and in all cases the average pre-stimulus baseline was subtracted from the post-stimulus epoch. The duration of this baseline was on average 193.75 ms (SD =28.16, range = 100 -250). The average post-stimulus epoch duration is shown in Figure 4C.

All studies reported using at least one method to remove or correct for artifacts. Artifact rejection procedures included rejecting epochs by visual inspection or using a fixed amplitude threshold. Artifact correction procedures included removing components after independent component analysis (ICA) or using various algorithms to remove EOG activity from the data. Some studies reported using additional filters to remove artifacts without providing further details. The percentage of studies using each of the main procedures is shown in Figure 4A. Automatic rejection using a fixed threshold was the most used method and the average rejection threshold is shown in Figure 4B. When using an artifact rejection procedure, 50% of studies reported the average number of epochs removed. On average, 11.34% of trials were removed (SD = 6.04, range = 1.34 - 29%).

**Fig. 4.**
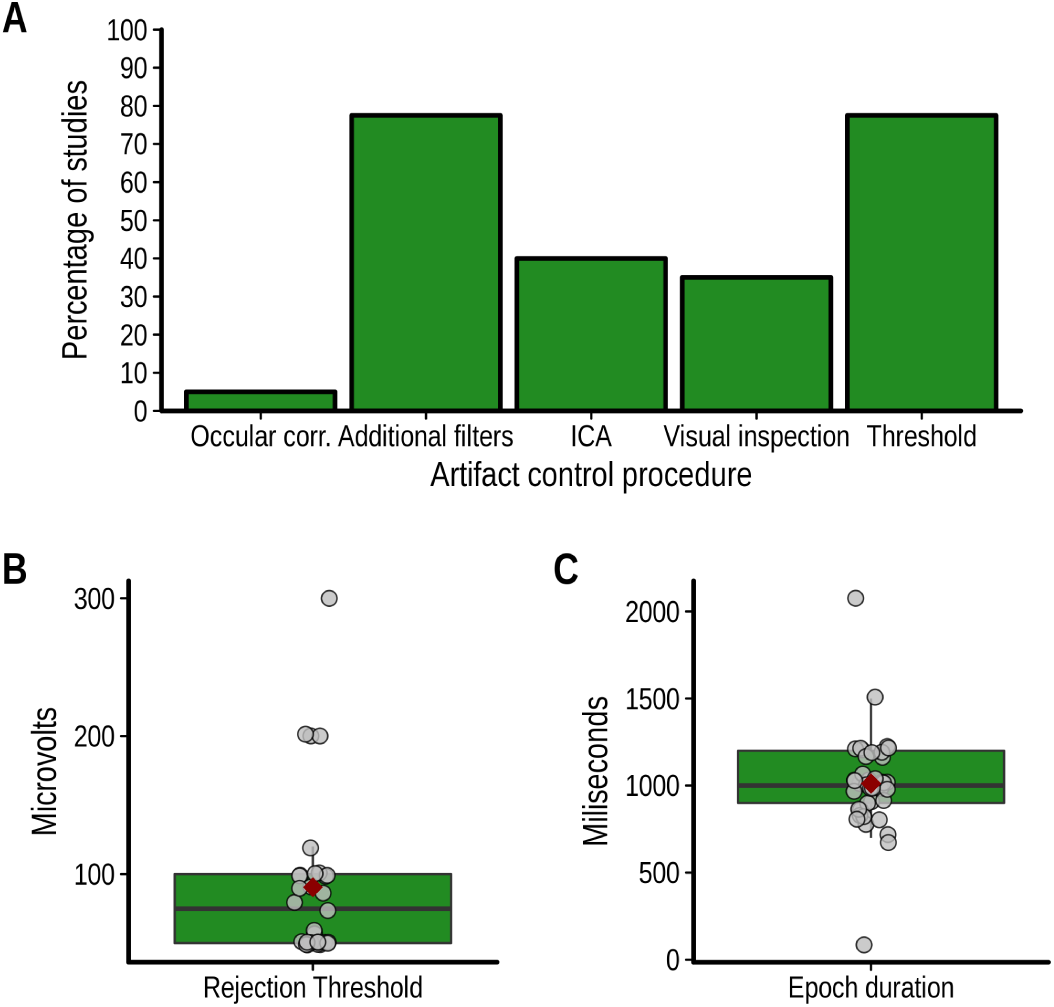
(**A**) Percentage of studies reviewed using each type of artifact control procedure identified, (**B**) value of the rejection threshold if used and (**C**) duration of epochs.

### ERP analyses

***ERP selection and measurement***. The average number of components analysed is shown in Figure 5C. In most cases, the choice of ERP components to analyse was based on previous studies (72.5%) while other studies chose components based on the inspection of the grand average waveform (17.5%), used another analysis to select the components of interest (2.5%) or did not justify their selection of components (7.5%). As shown in Figure 5A, the most widely analysed ERP components were the N1, N2, P2, P3 and the LPP. Note that in some cases, slightly different names were used for these components (e.g. P320 instead of P3). Furthermore, some studies also performed several analyses on the same component (e.g. early and late LPP). See Table S1 for the names and all components analysed in each study.

**Fig. 5.**
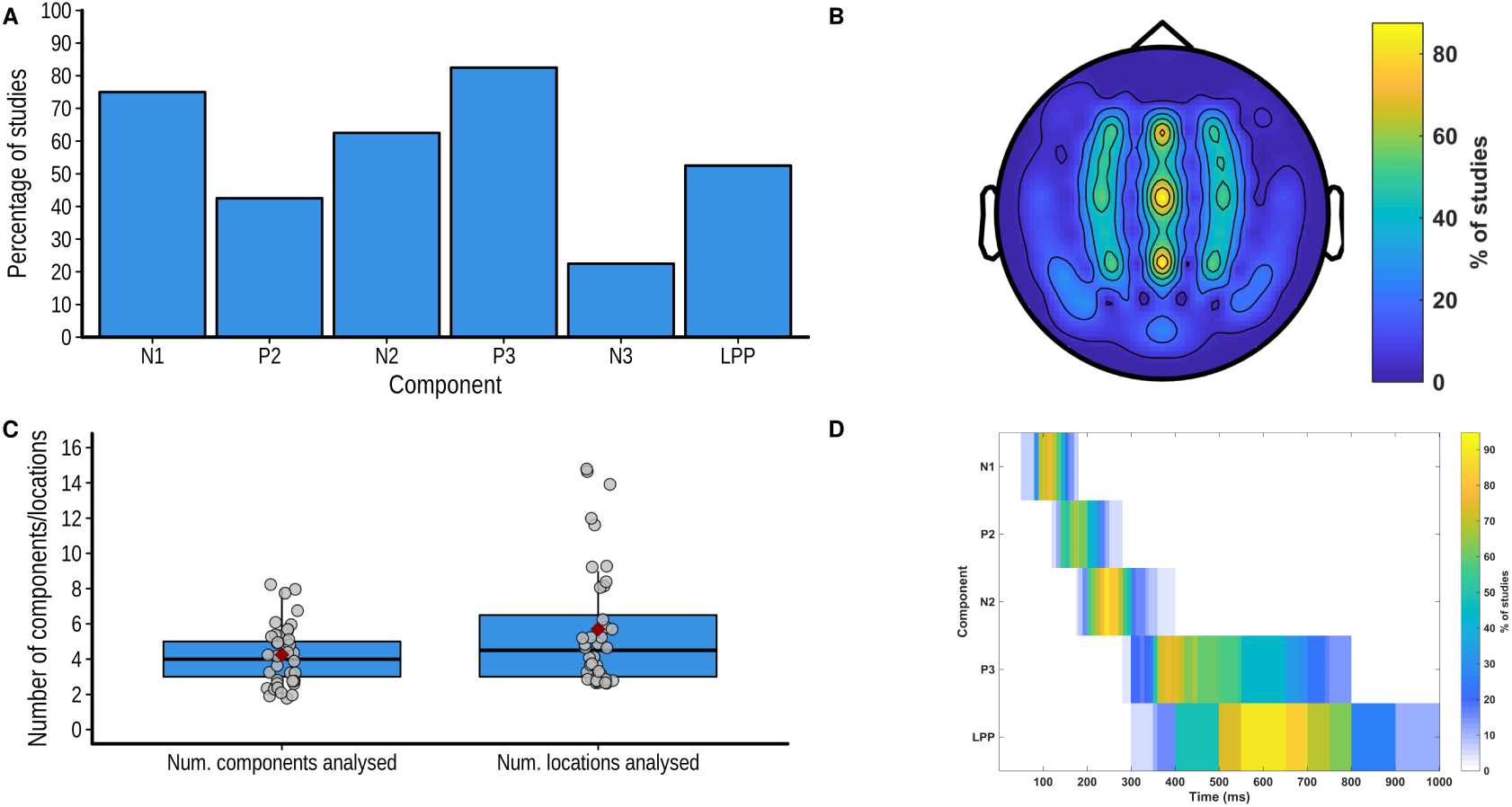
(**A**) Number of components and locations analyzed in each study. (**B**) Percentage of studies analyzing each of the main component. (**C**) Percentage of studies analyzing each scalp location. (**D**) Percentage of studies analyzing each component analyzing each time-point in the post-stimulus window.

The average number of locations analysed is shown in Figure 5C and the percentage of studies analysing each scalp location is shown on a 64 electrode montage in Figure 5B. A minority of studies provided a justification for the choice of locations to analyse (40%) and in most cases this choice was based on previous studies (25%). Almost all studies quantified the ERP components using the mean amplitude within a time window (80%) while other studies used the peak amplitude within a time window (7.5%) or point by point analyses (i.e., performing analyses in small time windows covering the whole ERP epoch; 7.5%). One study used peak amplitude or mean amplitude depending on the component (5%). The choice of the time window to analyse was either based on visual inspection (47.5%), previous studies (22.5%), both inspection an previous studies (previous studies (5%) the data itself (i.e circular analyses, 15%) or not justified (10%). The percentage of studies analysing each time point in the post-stimulus window for each component is shown in Figure 5D.

***ERP statistical analyses***. Almost all studies used factorial analyses of variance (ANOVA) to assess the statistical significance of the experimental factors on the ERPs (95%). One study used the analysis of covariance (ANCOVA) and another used the non-parametric Kruskal-Wallis test. Since the few studies using point-by-point analyses (7.5%) performed a large number of ANOVAs compared to the rest of the studies (on average 130 ANOVAs), these studies were not considered in the following description of the factorial analyses. Another study performing 54 ANOVAs was also considered an outlier and was not included in the descriptive statistics. The average number of ANOVAs performed in the remaining studies (N = 35) is shown if Figure 6. These ANOVAs had on average 3.2 factors (SD = 0.72, range = 2–5) and 19.6 cells (SD = 15.97, range = 4–60). To assess how many statistical tests these ANOVAs represent, the number of main effects and potential interactions was multiplied by the number of ANOVAs (see Luck & Gaspelin (2017)). The total number of tests in ANOVAs is shown in Figure 6. No studies corrected the significance threshold for the total number of ANOVAs performed. However, some studies (60%) corrected the significance threshold when performing post-hoc comparisons using the Bonferroni (42.5%), Tukey (10%), Scheffe correction (5%) or FDR (2.5%) correction. Several studies used the Greenhouse-Geisser correction when performing repeated-measure analyses (52.5%).

**Fig. 6.**
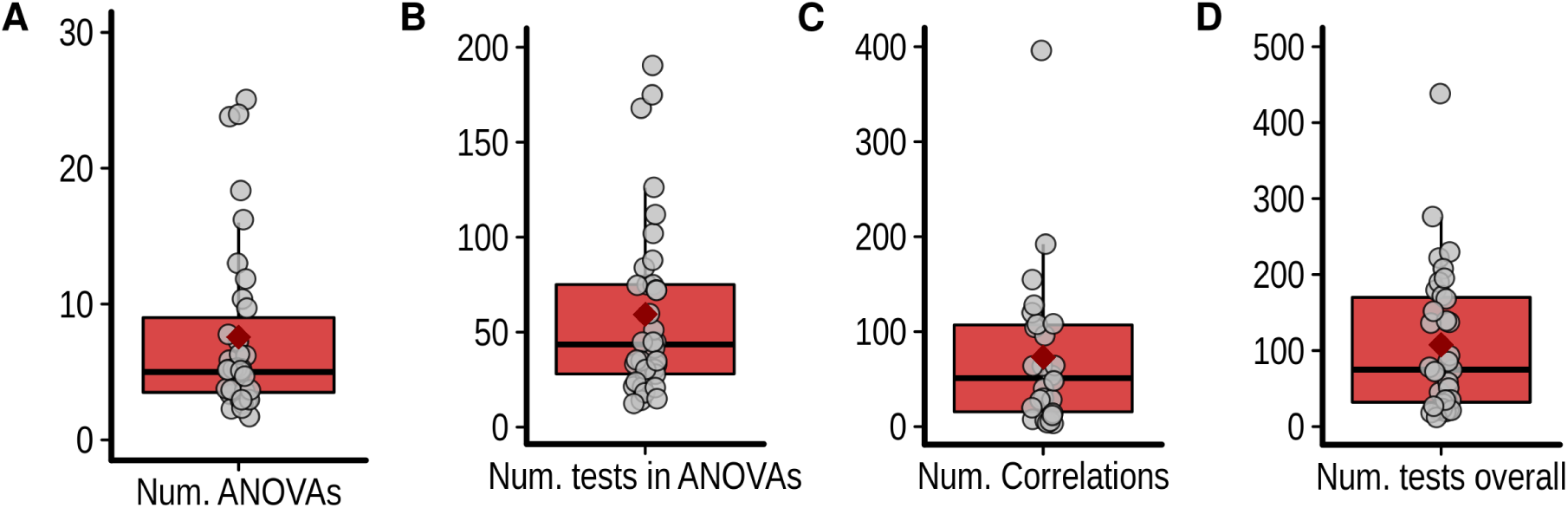
(**A**) Number of ANOVAs performed, (**B**) total number of statistical tests performed using ANOVAs, (**C**) number of correlations and (**D**) overall number of statistical tests performed.

65% of studies performed correlations between ERP data and other variables in addition to the factorial analyses. These correlations either used the difference between the ERP amplitude in two conditions (32.5%), the mean amplitude in a particular condition (27.5%), both mean amplitude and peak amplitude (2.5%) or mean amplitude and peak latency (2.5%) and were often performed in multiple time windows and scalp locations. The average number of correlations per study is shown in Figure 6. 10 % of studies corrected the significance threshold to control for the possibility of a type 1 error in correlation analyses.

### Results reporting

92.5% of studies reported mainly or exclusively significant results. When reporting the results from factorial analyses, 50% of studies did not report any estimate of effect size while the rest reported the partial eta-squared (45%), Cohen’s d (2.5%) or both (2.5‘%).

All but one study plotted the ERP data. 32.5% plotted only the time-course of the ERP response while 65% plotted both the time course and scalp maps at particular time points. In most cases, the locations and time points plotted were chosen because they were thought to be representative of the results (57.5%) while other studies plotted all locations analyzed (25%) or only the locations showing significant effects (15%). On the time course plots, only one study plotted the error intervals. In addition to the time course and scalp maps, 22.5% of studies also reported the ERP amplitudes in a table and 42.5% in bar graphs.

## Meta-analysis of the effect of pain observation

The results from the meta-analysis of the effect of the observation of pain stimulation on the components and locations that were the most frequently analysed are shown in Figure 7 and 8. Forest plots for each component at each location are shown in Figures S1-S8.

**Fig. 7.**
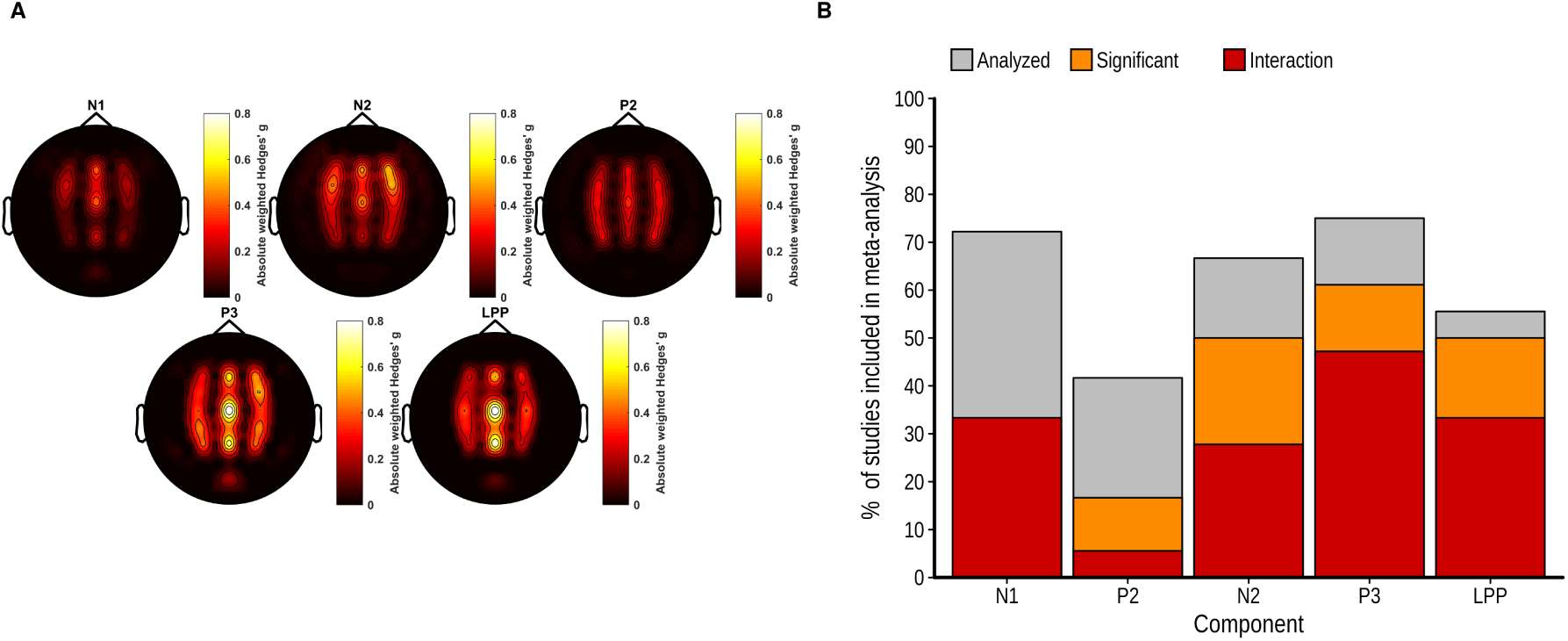
(**A**) Scalp map of the average Hedges’g weighted by the number of studies reporting results at this electrode for the main effect of pain observation on each electrode and component. (**B**) Bar graph showing the proportion of studies included in the meta-analysis analyzing each component, reporting a significant main effect of pain and a significant interaction between this effect and another factor.

**Fig. 8.**
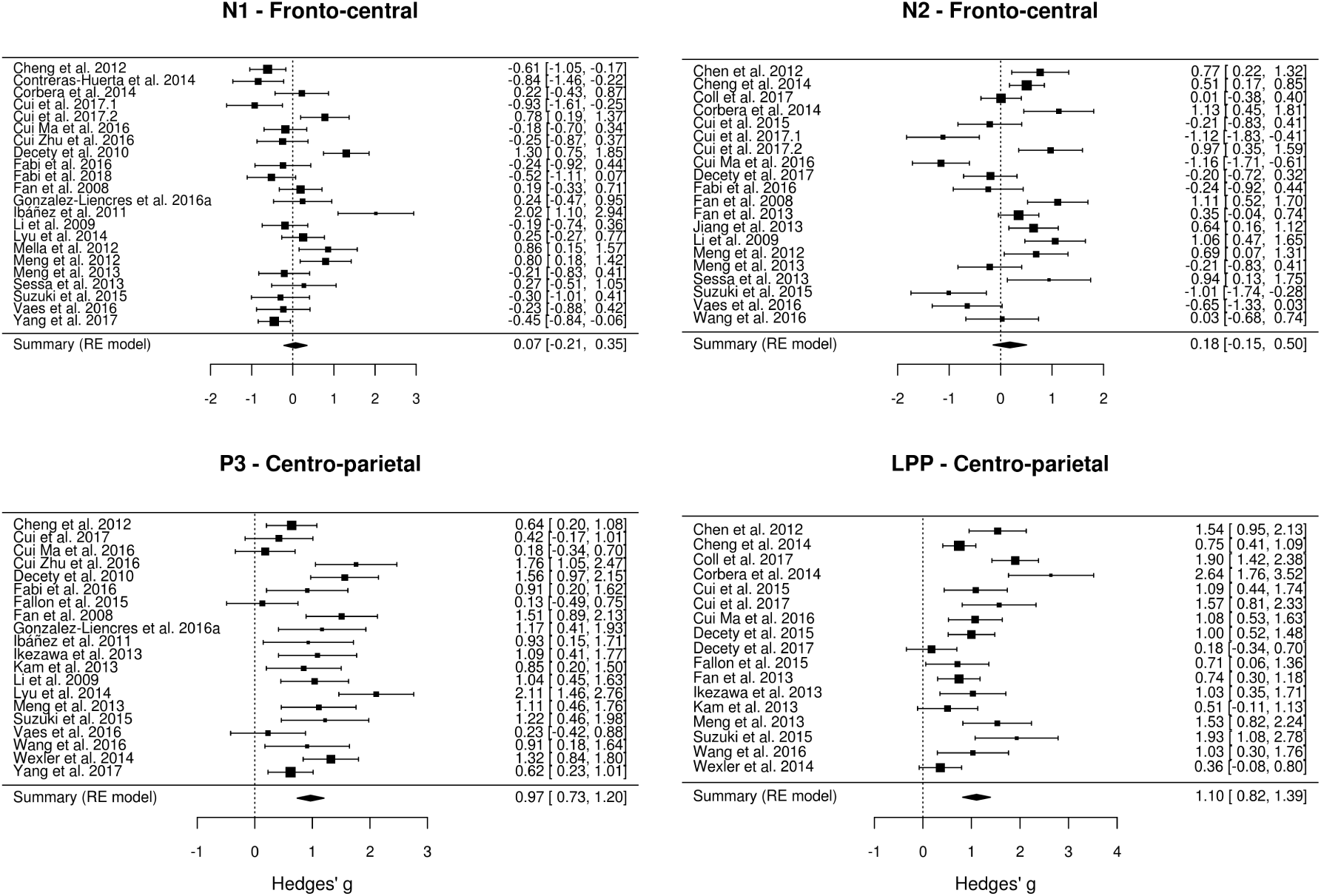
Forest plots showing the effect size and 95% confidence interval for the effect of pain observation on the ERP amplitude at the frontal electrodes fro the N1, N2, P3 and LPP components. Positive effects indicate higher (more positive) amplitudes for pain pictures and negative effects indicate lower (more negative) amplitudes for the pain pictures.

***N1 component***. The effect of pain observation on the N1 component is maximal at frontal electrodes (Figure 7A). Although a minority of studies measuring this component found a significant effect of pain observation, several studies found a significant interaction between the effect of pain observation and another experimental factor at any electrode site. The random-effect meta-analytic model fitted on the effect of pain on the N1 component at fronto-central sites collected from 22 studies suggests the presence of a high heterogeneity in the effect sizes (Q = 94.67, df =21, *p* < 0.001; I^2^ = 79.37%). Overall, the random-effect model indicated that of pain observation on the N1 component at fronto-central sites is not significant (*g* = 0.07, *k* = 22, *p* = 0.644, 95% CI: -0.21:0.35).

***N2 component***. Similarly to the N1 component, the effect of pain observation on the N2 component is also maximal at frontal electrodes (Figure 7A). Approximately 50% of studies included in the meta-analysis found a significant effect and 27.78% found a significant interaction between the effect of pain and another experimental factor at any electrode site. At the fronto-central sites, there is evidence for significant heterogeneity across the 20 studies(Q = 108.46, df =19, *p* < 0.001; I^2^ = 85.13%). Interestingly, the direction of the significant effects is highly heterogeneous, with a similar number of studies finding a significant increase or decrease in amplitude during pain observation. This led to a non-significant overall effect of pain observation on the N2 component at fronto-central sites. (g = 0.18, *k* = 20, p= = 0.283; I^2^ = 85.13, 95% CI: −0.15:0.5).

***P3 component***. While the effects of pain observation on the P3 component is distributed across the scalp, the effect is maximal at centro-parietal sites. Most studies measuring this component found a significant effect and a significant interaction between the effect of pain and another experimental factor. Although the effect sizes for the centro-parietal P3 component across the 20 studies are considerably less heterogeneous than those found for the early components, there is still significant heterogeneity across studies (Q = 55.78, df = 19, *p* < 0.001; I^2^ = 66.33). All studies found that pain observation led to a positive shift in P3 amplitude and the overall effect is large and significant (*g* = 0.97, *k* = 20, *p* < 0.001, 95% CI: 0.73:1.2).

***LPP component***. The effect of pain observation on the LPP component is strongest at centro-parietal sites. As for the P3 component, most studies measuring this component found a significant effect of pain observation and 33.33% found a significant interaction between the effect of pain and another experimental factor. At the centro-parietal location, there is significant heterogeneity across the 17 studies (Q = 63.62, df = 16, *p* < 0.001; I^2^ = 76.39%), despite the fact that pain observation led to a positive shift in LPP amplitude for all studies. The overall effect of pain observation on the LPP is large and significant (*g* = 1.1, *k* = 17, *p* < 0.001, 95% CI: 0.8195:1.39).

***Publication bias and excess significance***. Funnel plots illustrating the effect size for each study as a function of study precision are shown in Figure 9. It should be noted that the low variance in precision (due to most studies having a similar sample size) limits the interpretation of these figures. Nevertheless, the funnel plots show that the effect sizes were roughly symmetrically distributed across the summary effect size, suggesting the absence of publication bias. This observation was formally tested using linear regressions to asses the relationship between the magnitude of the effect size and the precision (standard error) of each study. This procedure revealed non-significant relationships between effect size and precision for the N1 component at fronto-central sites (t(20) = 1.84, *p* = 0.08), the N2 component at fronto-central sites (t(18) = −0.76, *p* = 0.45) and the P3 component at centro-parietal sites (t(18) = 1.46, *p* = 0.16). A significant negative relationship between effect sizes and precision is present for the LPP component at centro-parietal sites (b = 3.91, t(15) = 2.32, *p* = 0.03). This suggests that the large effect sizes found for the LPP component in some studies with a smaller sample size are likely inflated.

**Fig. 9.**
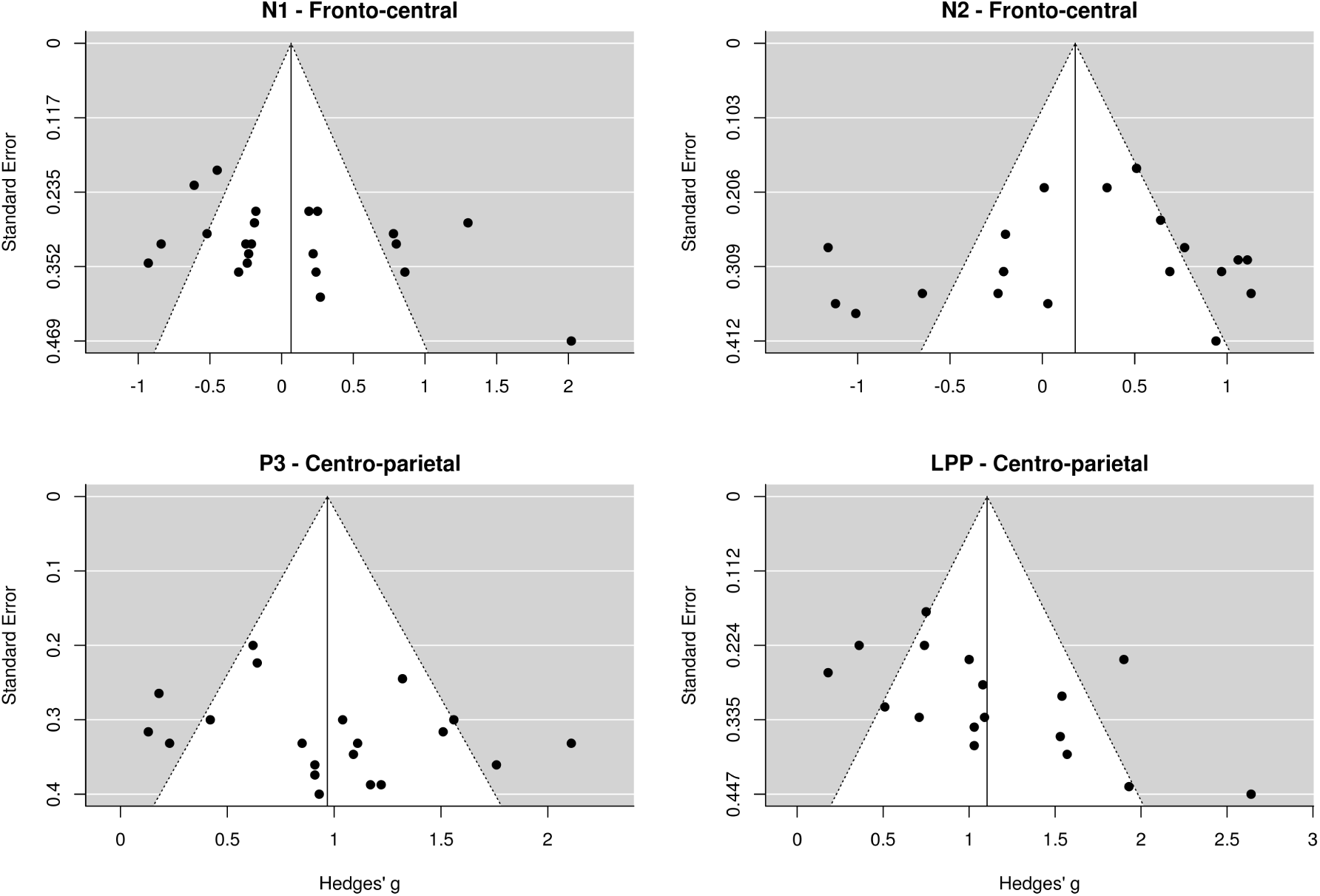
Funnel plots showing the distribution of studies as a function of effect size and precision (standard error). The white area shows the 95% confidence interval around the summary effect size for each degree of precision.

Exact binomial tests assessing the presence of excess significance suggest that the number of significant effects found for the N1 and N2 component is significantly higher than expected given the power of each study and the summary effect size (*p* < 0.001 for both). The proportion of significant effects is however similar to the expected number for the P3 and LPP components (*p* > 0.9 for both).

## Discussion

The meta-analysis of 36 studies investigating pain empathy using ERPs suggests that the observation of pain in others does not reliably modulates the early N1 and N2 components often measured in these studies. However, large and reliable effects were found for the later P3 and LPP components.

These findings challenge a popular model arguing that the time-course of the ERP to the observation of pain in others is characterised by an early frontal “affective sharing response” followed by a centro-parietal “late cognitive reappraisal” of the stimulus (Decety et al., 2010; Fan and Han, 2008). Indeed, the current results suggest that the observation of pain in others does not lead to a reliable modulation of early frontal ERP components thus undermining the idea that they are associated with automatic emotion sharing. It is possible that the vicarious pain stimuli used in the studies reviewed led to a modulation of early posterior components as is generally observed for emotional stimuli (Schupp et al., 2003b,a). Unfortunately, almost none of the studies reviewed measured early responses at posterior sites, thus preventing the comparison between ERP components that are claimed to reflect empathic processes and the components usually observed in response to emotional stimuli.

The results of the meta-analysis are also compatible with a wealth ERP research showing that emotional stimuli reliably modulate the later centro-parietal components (Hajcak et al., 2010; Schupp et al., 2003a). However, since this modulation is commonly observed in response to many types of emotional stimuli, the idea that it represents an empathic response in the context of pain empathy and not a general aversive/regulatory response remains to be established. Indeed, none of the study reviewed compared the ERP response to vicarious pain stimuli to non-social emotional stimuli to assess the specificity and uniqueness of the processes indexed (Happé et al., 2017). It seems imperative for future studies using ERP to investigate pain empathy to carefully evaluate the validity, reliability and specificity of ERP responses to pain in others.

Despite the lack of reliability of the early frontal effects observed here, a considerable proportion of the studies reviewed reports significant effects of pain observation on the early frontal N1 and N2 components. This apparent contradiction could be explained by the methodological issues underlined in the systematic review of methodological practices indicating that there was considerable variability in the quantification and statistical analysis of the ERP data. Indeed, while most studies used the mean amplitude between fixed-latencies to quantify ERPs, the time-windows and the electrode locations used for this measure often varied considerably across studies without a clear rationale underlying this choice. The problems of analytical flexibility, sometimes called researcher’s degrees of freedom, have already been discussed elsewhere (Luck & Gaspelin, 2017; Simmons et al., 2011). However, the current study suggests that previous investigations of pain empathy using ERPs might be compromised by this practice.

This analytical flexibility was often combined with a shotgun analytical approach in which a high number of statistical analyses were performed on several locations and time-windows and any significant effect was interpreted as meaningful. On average, the studies reviewed here performed X statistical tests on ERP data and none corrected the significance threshold to reduce the risk of a false positive finding. This is problematic since such a high number of statistical tests leads to a high probability that several significant results are in fact false positives (Kilner, 2013; Luck & Gaspelin, 2017). To diminish this risk, researchers can use analytical techniques that can take into account the spatial and temporal distribution of ERP data to reduce the number of comparisons or to adequately control for them (Groppe et al., 2011; Pernet et al., 2015; Maris & Oostenveld, 2007).

Alternatively, researchers can restrict their analyses to scalp regions and time windows for which an effect was predicted. However, a worrying result of this study is the fact that in several cases the analysis and interpretation of the data were not constrained by the researchers’ predictions since no clear hypotheses were formulated. This suggests that despite using the confirmatory analytical approach of null hypothesis significance testing, most of ERP research in the field of pain empathy is exploratory (Wagenmakers et al., 2012), even after 10 years and 40 studies. A solution to this issue would be to require researchers to clearly formulate their research hypotheses and label as exploratory the results of analyses that were not predicted. Ideally, researchers could pre-register their hypotheses online or publish using the pre-registered report format to establish the analysis plan before collecting data (Munafo et al., 2017).

Another striking observation permitted by this review is the lack of comprehensive reporting for the results from the statistical analysis of ERP components. Indeed, it was found that the vast majority of studies only reported significant results form a large number of factorial analyses. Therefore, in addition to a potential publication bias (Rosenthal, 1979), the ERP studies reviewed here also show a within-study reporting bias according to which analyses leading to non-significant results are less likely to be reported. Putting aside the fact that negative results can sometimes be informative if statistical power is high enough (Greenwald, 1975), the main consequence of this practice is that any attempt to meta-analytically summarize the results of such studies will be difficult, inevitably biased and of questionable usefulness (Moran et al., 2017). Therefore, it should be noted that the effect size calculated in the meta-analysis performed in this report are probably inflated.

In several cases, this practice was justified by the necessity to provide a concise and brief report of the results. The short term solution to this issue is to make it mandatory for authors to appropriately report all the results of all statistical analyses performed on ERP components in the text or in supplementary materials. It would also be beneficial for the field of ERP research to adopt a standard reporting procedure that would enable the automatic extraction of results from published articles and facilitate meta-analyses and large-scale automated summaries of all published studies. For example, the field of functional magnetic resonance imaging research has taken advantage of the standard reporting of activation coordinates in tables to produce automated meta-analytical tools (Yarkoni et al., 2011). A more preferable long term solution would be to encourage the sharing of ERP data in online repository which would allow the re-analysis and meta-analysis of large datasets and a quick and efficient assessment of the evidence for specific effects (Poldrack & Gorgolewski, 2014).

A related issue to the incomplete reporting of results was incomplete and unclear reporting of several important methodological details. For example, less than half of the studies reported the number of trials left after artifact rejection procedures, meaning that it was unknown how many trials were included in the analyses in a majority of cases. To avoid omitting to report such crucial information, researchers should refer to published guidelines for reporting of EEG experiments (Keil et al., 2014; Moran et al., 2017; Picton et al., 2000) and reviewers should enforce these guidelines in all relevant cases.

### Limitations

Due to the issues discussed above, the results from the meta-analysis are most likely biased. This means that while they can be used to roughly guide future investigations, the precise value of the effect sizes should not be taken at face value.

The high heterogeneity in the experimental designs and analytical approaches as well as partial reporting of results in the studies analysed prevented an analysis of factors modulating the ERPs to vicarious pain. It is therefore possible that the lack of effect found for the early components is due to a moderation by other factors that could not be assessed. The improvement of methods and reporting in future studies should allow a more comprehensive analysis of factors potentially modulating the effects.

The methodological issues highlighted in this report are not specific to the domain of pain empathy and the present observations could potentially be generalised to many other fields of research in cognitive neuroscience (e.g. Hobson & Bishop (2017), for similar observations on EEG studies of action observation). Furthermore, while several issues were found to be prevalent in the studies reviewed, the scientific quality and usefulness of all papers cited in this report should be assessed on an individual basis.

Finally, the solutions proposed to the issues raised are not exhaustive, nor can they be applied indiscriminately to all ERP research.

## Conclusion

In conclusion, this study provides meta-analytic evidence for a robust modulation of later, but not early ERP components during pain observation. Furthermore, it suggests current framework used in pain empathy research using ERPs the investigation of the empathic response to pain in others using ERPs is undermined by several methodological problems that raise doubts regarding the reliability, validity and overall usefulness of this research. Researchers in the field should take into account the methodological issues raised here when designing and reviewing ERP experiments. This is of critical importance if this paradigm is to be used to draw conclusions on socio-emotional functioning in different clinical populations.

## Acknowledgements

MP Coll is funded by a fellowship from the Canadian Institutes of Health Research and was funded by a fellowship from the Fonds de Recherche Quebec - Sante during this work. The author declares no conflict of interest.

